# Helminth egg derivatives as pro-regenerative immunotherapies

**DOI:** 10.1101/2022.05.02.490277

**Authors:** David R. Maestas, Liam Chung, Jin Han, Xiaokun Wang, Sven D. Sommerfeld, Erika Moore, Helen Hieu Nguyen, Joscelyn C. Mejías, Alexis N. Peña, Hong Zhang, Joshua S. T. Hooks, Alexander F. Chin, James I. Andorko, Cindy Berlicke, Kavita Krishnan, Younghwan Choi, Amy E. Anderson, Ronak Mahatme, Christopher Mejia, Marie Eric, JiWon Woo, Sudipto Ganguly, Donald J. Zack, Franck Housseau, Drew M. Pardoll, Jennifer H. Elisseeff

## Abstract

The immune system is increasingly recognized as an important regulator of tissue repair. We developed a regenerative immunotherapy from the helminth *Schistosoma mansoni* soluble egg antigen (SEA) to stimulate production of interleukin (IL)-4 and other type 2-associated cytokines without negative infection-related sequelae. The regenerative SEA (rSEA) applied to a murine muscle injury induced accumulation of IL-4 expressing T helper cells, eosinophils, and regulatory T cells, and decreased expression of IL-17A in gamma delta (γδ) T cells, resulting in improved repair and decreased fibrosis. Encapsulation and controlled release of rSEA in a hydrogel further enhanced type 2 immunity and larger volumes of tissue repair. The broad regenerative capacity of rSEA was validated in articular joint and corneal injury models. These results introduce a new regenerative immunotherapy approach using natural helminth-derivatives.

**One-Sentence Summary:** Helminth-derived soluble egg antigen regenerative immunotherapies promote tissue repair in multiple injury models.

## Main Text

Regenerative medicine therapies traditionally aim to promote new tissue development by either stimulating endogenous resident stem cells or through the delivery of exogenous stem cells (*1*). Biomaterials and biological signals in the form of growth factors or drugs have also been designed and introduced to further enhance cell activity and tissue development. Unfortunately, the efficacy of these therapies often decreases as they move to testing in larger species and clinical impact remains limited (*2*). The immune system is now recognized as an important contributor to tissue repair through production of cytokines that stimulate stem cells and creation of a microenvironment that supports new tissue development (*3*). Multiple immune cells and immunological active stromal cells contribute to the microenvironment in ways that can inhibit or promote tissue repair. New regenerative medicine therapies that target the immune system may provide an alternative strategy to stimulate tissue repair by creating a pro-regenerative tissue immune microenvironment. We found previously that acceleration of muscle wound repair by application of extracellular matrix (ECM)-derived biological scaffolds was dependent on induction of T helper type 2 (T_H_2) cell responses, characterized by the mechanistic target of rapamycin-2 (mTORC2) pathway activation and production of interleukin (IL)-4, and other groups have reported that IL-4 producing eosinophils are also required (*4, 5*). This discovery prompted us to evaluate the potential for enhanced wound healing by helminthic products, since parasitic worm infections are widely regarded to be among the strongest inducers of T_H_2 and type 2 immunity.

Parasitic worm infections affect billions of people worldwide, primarily in tropical developing regions, where they can cause morbidity and mortality (*6-8*). One such parasite, *Schistosoma mansoni* (*S. mansoni*), is a helminth that can infect and can cause severe damage to the liver and hepatic blood vasculature due in part to the immune responses against the eggs lodged in hepatic sinusoids. These responses to xenogeneic foreign bodies and their respective secretome lead to granuloma formation, and if left unresolved can result in fibrosis (*9-12*). While *S. mansoni* and other helminth infections can cause a variety of disease states, they also appear to promote a number of desirable features. For example, some helminth infections reduce the incidence of allergies and are associated with beneficial alterations in the microbiome (*13, 14*). Infection has also been associated with decreased symptom severity in auto-immune disease and even Alzheimer’s disease symptoms, suggesting helminths may induce beneficial immunomodulation (*15-17*). More directly connecting helminth responses to tissue repair, Nusse *et al*. demonstrated that a *Heligmosonoides polygyrus* infection triggers epithelial stem cell niche reprogramming, suggesting a connection between helminths and tissue repair that is modulated by the immune response to the worm and its egg secretome (*18, 19*). Finally, helminth infection induces IL-4 receptor (IL-4R) signaling that downregulates IL-17A production to benefit rapid tissue repair after the initial infection (*20*).

Type 2 immunity is characterized by the influx of T_H_2 cells, eosinophils, type 2 innate lymphoid cells (ILC2), alternatively activated macrophages, and type 2 associated cytokines IL-4, IL-5, IL-9, and IL-13 (*18, 21*). Previous studies found that the egg secretome components of *S. mansoni* and the soluble egg antigen (SEA) can stimulate type 2 immunity, recapitulating in part the immune response to infection. Evidence suggest that multiple components of SEA can contribute to the type 2 immune responses, in particular the glycoproteins Omega-1, IPSE/alpha-1(IL-4 inducing principle of *S. mansoni* eggs), and the Lewis^X^ containing glycan LNFP III (*22-24*).

While the type 2 immune response is historically considered a protective response against parasitic worms, new perspectives suggest that helminths may be co-opting this immune response to repair the damage induced during infection by agonizing type 2 cytokines. For example, IL-4 secreted by eosinophils and T_H_2 cells can promote activation of other cell types required for skeletal muscle repair (*4, 5*). In the central nervous system, IL-4 secreting T_H_2 cells promote repair in the retina and spinal cord and IL-4 levels are positively associated with learning in Alzheimer’s disease animal models (*25-27*). Although helminths drive type 2 responses, it is unknown whether helminth secretory agents can be formulated or engineered as safe and effective regenerative immunotherapies without the pathologies coincident with parasitic infections such as fibrosis and hepatic damage (*28-30*). In this work, we developed a formulation of SEA from *S. mansoni* eggs that induces a type 2 immune response and promotes tissue repair while decreasing signatures of pathological inflammation and fibrosis, opening the door to a new approach to regenerative immunotherapy.

## Results

### Development of a type 2 immunotherapy from S. mansoni egg antigens

The soluble egg antigen (SEA) extract from *S. mansoni* is composed of hundreds to thousands of proteins, glycoproteins, and lipids depending on the extraction protocol, and is well recognized in its ability to stimulate a type 2 immune response (*31-34*). We first sought to determine if SEA could efficiently stimulate a type 2 immune response that would promote tissue repair without deleterious inflammation or fibrosis. *S. mansoni* eggs isolated from infected mice were mechanically disrupted and ultracentrifuged to remove insoluble components to isolate SEA as described by Boros, et al. (**fig. S1a-c**) (*28*). We first evaluated the immune response to SEA using an *in vitro* screen with splenocytes from murine C57BL/6 wild type and IL4-eGFP reporter (4get) mice. Addition of SEA to the splenocyte culture resulted in increased expression of IL4-eGFP by CD4 T cells comparable to IL-4 (**fig. S2a-b**), however there was also a modest increase in CD8^+^ T cells expressing IFN-γ as measured by cytokine staining in C57BL/6 splenocyte cultures (**fig. S3a-c**). To further test in a wound healing model, we created a volumetric muscle loss wound (VML) to evaluate the repair capacity of SEA. Application of a single dose of SEA to the VML at the time of injury resulted in a significant increase in *Il4* gene expression in the muscle tissue 1-week post-surgery (**fig. S4**). However, similar to the *in vitro* studies, there was also a significant increase in *Il1b* expression and increased expression of *Ifng* and *Tnfa* in the muscle tissue with SEA treatment compared to saline treated control injury (**fig. S4**).

To determine if SEA could be formulated to stimulate a dedicated type 2 regenerative immune response without increased interferon-dependent pathological inflammation, we modified the SEA isolation to extract specific layers of the ultracentrifuged egg supernatant. In particular, we further purified the SEA by isolating the lower density soluble fraction and combined it with the lipid portion, as recent lipidomic research reported that *S. mansoni* worms and eggs contain pro-resolving mediators which can promote wound healing (*35, 36*). Protein gel electrophoresis confirmed that the modified isolation protocol produced a more purified SEA formulation compared to traditional protocols (**fig. S5a**). The *in vitro* 4get splenocyte screening assay confirmed IL-4 stimulation along with cell proliferation with the modified SEA formulation (**fig. S5b, S6a-b**) that was consistently generated from multiple batches (**fig. S6c, d**).

Application of the modified SEA formulation to the VML injury model resulted in a significant increase in *Il4* gene expression in the muscle tissue without a concomitant increase in *Ifng* and *Il1b* expression (**Fig. 1a, fig. S7**). Instead, expression of these inflammatory genes decreased with the modified SEA treatment compared to standard SEA and saline treated injuries one-week post-surgery. Bulk RNA sequencing confirmed significant differences between classical SEA and the modified SEA formulation (**fig. S7**). This formulation was then used for all subsequent studies and is referred to as regenerative SEA (rSEA). We then evaluated the different delivery modalities for the rSEA formulation by comparing local and systemic (intraperitoneal, IP) injection. Both local delivery in the VML wound and systemic IP injection of rSEA increased IL-4 expression in the muscle tissue one week after injury (**fig. S8a-b**). However, further examination of specific immune subsets found that local rSEA treatment significantly increased both IL-4 expressing T cells and eosinophils more than double compared to systemic delivery (**fig. S8b**) so we used local treatment for all subsequent studies.

**Fig. 1.**
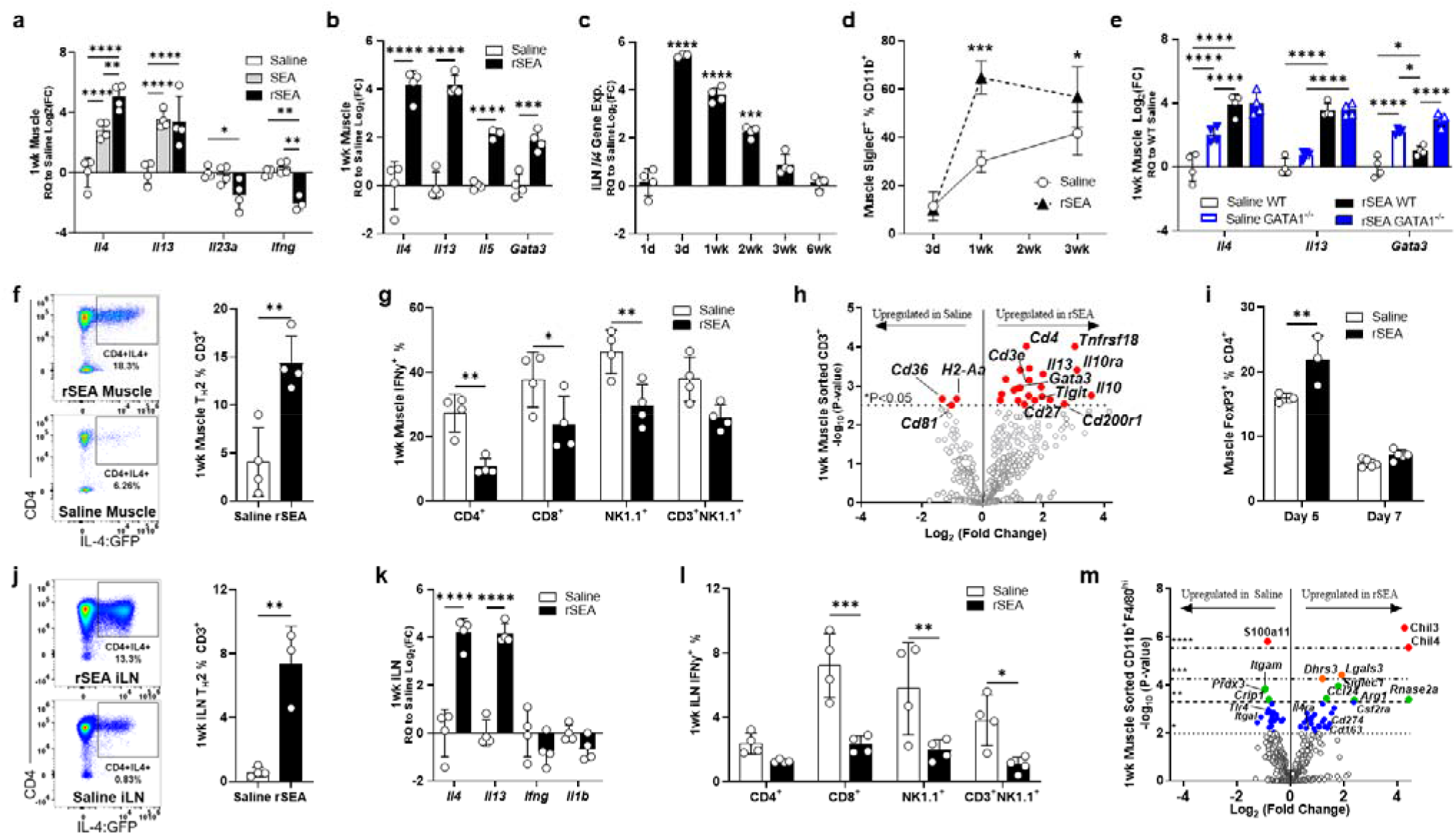
rSEA treatment promotes a pro-regenerative type 2 immune microenvironment after muscle injury. Mice had a partial quadriceps resection to create volumetric muscle loss (VML) injury and received the indicated treatments administered locally at the time of resection. **a**,**b**. Muscle tissue expression of selected T_H_2 and T_H_1 genes assessed by qRT-PCR 1-week post-VML and treatment with saline, unfractionated soluble SEA extract and fractionated SEA formulation (regenerative SEA, rSEA). **c**. *Il4* gene expression of iLNs from VML injured mice at different time points after rSEA treatment. **d**. Flow cytometric quantitation of post-injury muscle eosinophils, defined by co-expression of Siglec F and CD11b, over time with rSEA treatment. **e**. Muscle expression of selected type 2 genes after saline vs rSEA treatment of Wild Type (WT) and GATA1 KO mice. **f**. Flow cytometry of muscle CD3^+^CD4^+^ cells from IL4-reporter (4get) mice treated with either saline or rSEA. **g**. Muscle IFNγ^+^ production by various immune cell types, assayed by intracellular cytokine staining. **h**. Gene expression profile of sorted CD3^+^ T cells from muscle 1-week post-injury, comparing saline and rSEA. **i**. CD4^+^Foxp3^+^ (Tregs) in the muscle 5 and 7 days after treatment with either saline or rSEA. **j**. Flow plots of iLNs from 4get mice draining VML treated one week with saline or rSEA. **k**. iLN gene expression 1-week after muscle injury and treatment. **l**. iLN IFNγ production by various cell types assayed by ICS. **m**. Gene expression profiling of CD11b^+^F4/80^+Hi^ macrophages sorted from muscle one week after saline vs rSEA treatment. Statistical tests represent all biological replicates, and all experiments excluding (h, m) were replicated at least twice. Graphs show mean ± s.d. (a-g, i-l) and mean ± SEM, n = 3-5 (e). *\*P* < 0.05, *\*\*P* < 0.01, *\*\*\*P* < 0.001, *\*\*\*\*P* < 0.0001 by unpaired two-tailed Student’s t-test (f, j) and two-way ANOVA with Sidak’s multiple comparisons. Dashed lines (h, m) represent -Log_10_(Adj. P-value) determined by nCounter software.

### rSEA treatment increases IL4-expressing eosinophils, T_H_2 T cells, and regulatory T cells in a volumetric muscle loss injury

Since type 2 immune signals are critical for muscle injury repair, we evaluated the kinetics and cellular sources of IL-4 after rSEA treatment. A single treatment of rSEA at the time of injury increased *Il4* gene expression in the muscle tissue 20- to 30-fold, relative to saline treated muscles, at one-week post-surgery (**Fig. 1b**). Treatment with rSEA also increased expression of other type 2 associated genes including *Il5, Il13*, and *Gata3* (**Fig. 1b**). Furthermore, the *Il4* gene expression in the draining inguinal lymph nodes increased, peaking at 3 to 7 days post-treatment and returning to the level of saline treated controls after 3-weeks (**Fig. 1c**), suggesting that a single rSEA treatment was able to modulate the tissue immune environment for weeks.

Eosinophils are well recognized as a source of IL-4 after muscle injury so we first evaluated the changes in these cell types after rSEA treatment using multiparametric flow cytometry. Eosinophils (CD11b^+^SiglecF^+^SSC^Hi^), which depend on type 2 immune cytokines and chemokines such as IL-5 and eotaxin, were the most abundant CD11b^+^ cell population in the muscle 1-week post rSEA treatment producing an 11-fold increase in the number of cells that remained significantly higher compared to saline treated controls at 3-weeks post-treatment (**Fig. 1d**). However, *Il4* gene expression in the muscle tissue still increased in ΔdblGATA (GATA1) knock-out mice, which lack eosinophils, suggesting that rSEA is stimulating other cell types in the muscle wound (**Fig. 1e**).

Previous studies with ECM biomaterials demonstrated that T_H_2 cells are critical for the pro-regenerative response to ECM biological scaffolds so we further examined the T cell response to rSEA (*4, 37, 38*). At 1-week post-treatment, CD3^+^ T cell numbers increased in the muscle wound with CD3^+^CD4^+^ T cells increasing over 7-fold in the muscle with rSEA treatment compared to saline treated controls. Treatment with rSEA significantly increased T_H_2 (CD3^+^CD4^+^GFP^+^(IL4^+^)) as a percentage of CD3^+^ cells in the muscle of 4get mice (**Fig. 1f**). Since the 4get mice report the transcriptional status of the IL-4 locus via GFP expression and have a BALB/c background, we further confirmed direct increases in IL-4 protein with intracellular cytokine staining (ICS) flow cytometry in C57BL/6 mice and found significant increases in T_H_2 cells (CD3^+^CD4^+^IL4^+^) with rSEA treatment (**fig. S8c**). There were also changes in the more inflammatory subsets of T cells with rSEA treatment including a significant decrease in the percentage of IFN-γ expressing CD4^+^ T cells, a decreased percentage of CD8^+^ T cells, and decreased percentage of IFN-γ^+^ natural killer and natural killer T cells (**Fig. 1g**) suggesting that rSEA does not induce negative inflammatory changes in addition to the pro-regenerative response.

We further characterized T cell gene expression changes with rSEA treatment by sorting CD3^+^ T cells from 1-week post-injury muscle and analyzing gene expression signatures with the NanoString multiplex system. There were 20 genes significantly upregulated and downregulated in T cells with rSEA treatment (**Fig. 1h, Table S1**). In particular, rSEA treatment of the muscle wound significantly upregulated T_H_2 associated gene signatures including *Il13, Gata3, and Cd4*. Expression of *Tnfrsf18, Ccr4, Il10*, and *Il10ra* also significantly increased in the T cells after rSEA treatment suggesting a role for regulatory T cells (T_regs_). Flow cytometry confirmed a significant increase in T_regs_ (CD3^+^CD4^+^Foxp3^+^) in the muscle with rSEA treatment compared to saline controls on days 5 and 7 post-injury (**Fig. 1i**), and higher T_reg_ percentages in the lymph nodes draining the muscle injury sites (**fig. S9a**). As T_regs_ are known to be integral to muscle healing (*39*), this further supports a regenerative phenotype induced by rSEA.

The draining inguinal lymph nodes (iLNs) also revealed type 2 immune stimulation after rSEA treatment including a significant increase in IL4^+^CD4^+^ T_H_2 cell percentage in 4get mice (**Fig. 1j**) and significant increases in T_H_2-associated gene expression signatures combined with decreased *Ifng* and *Il1b* at 1-week post-injury (**Fig. 1k**). Similar to the muscle, we further confirmed the iLN’s expression of IL-4 with ICS flow cytometry in C57BL/6 mice and found that rSEA induced T_H_2 cells (CD3^+^CD4^+^IL4^+^, **fig. S9b**). There was a significant decrease in the percentage of IFN-γ production in CD4^+^ T cells, a decreased percentage of CD8^+^ T cells, and decreased percentage of IFN-γ^+^ natural killer and natural killer T cells (**Fig. 1l**). While there were no differences in CD19^+^ B cells in the muscle 1-week post-treatment, we found significant changes in B cell percentages and phenotypes in the iLNs (**fig. S10a**). The number of CD19^+^B220^+^ B cells in the iLNs increased compared to saline controls and the percentage of CD19^+^B220^+^ B cells also increased in the iLNs (**fig. S10b**). In the *in vitro* splenocyte cultures, rSEA exposure significantly increased B cell numbers, suggesting a direct stimulatory role of rSEA on B cells in the regenerative immune response (**fig. S11**).

### rSEA induces alternatively activated macrophage gene expression in muscle wounds

Macrophages are another immune cell type that is central to tissue repair with alternatively activated macrophages associated with productive wound healing (*40*). The number of macrophages in the muscle tissue had no significant change with rSEA treatment (**fig. S12a, b**) and there were no differences in CD86 and CD206 expression (**fig. S12c**). To further evaluate the macrophages, we sorted CD11b^+^F4/80^Hi+^ myeloid cells from the muscle wound 1-week post-injury and utilized the NanoString Myeloid Codeset for gene expression analysis. rSEA treatment induced significant changes in macrophage expression of 57 out of the 770 genes tested (**Fig. 1m**). We found expression of 31 genes significantly increased with rSEA treatment related to metabolism, cell migration/recruitment, and cell activation including *Chil3, Arg1, Cd163, and Ccl24* (Eotaxin-2) that are correlated with non-inflammatory macrophages (**Fig. 1m**). Confirming expression of these genes with qRT-PCR, we found that rSEA treated muscle wounds significantly increased expression of *Chil3, Rnase2a*, and *Arg1* compared to controls (**fig. S12d**). The CD11b^+^F4/80^Hi+^ cells sorted from rSEA-treated muscle also significantly downregulated several genes associated with inflammation and complement activation genes such as *S100a11, Itgam, Itgal, Cxcl16*, and *C1qc* (**Fig. 1m, Table S2**). The most strongly downregulated gene, *Cysltr1*, encodes for cysteinyl leukotriene receptor-1, a potent mediator of allergic inflammation. In contrast to an active helminth infection, rSEA treatment resulted in downregulation of genes such as *Tlr2, Tlr4* and *Jun* in the sorted myeloid cells compared to saline treated controls. This suggests that the rSEA formulation can induce the regenerative components of the type 2 immune helminth response without the deleterious infection response component.

### rSEA stimulation of type 2 immunity correlates with increased muscle repair

IL-4 expressing eosinophils, T_H_2 cells and regulatory T cells are all associated with muscle repair after traumatic injury (*4, 5, 39, 41*). To assess whether rSEA stimulation of type 2 immunity also benefitted muscle repair, we assessed healing and fibrosis with histology and further expression analysis at early (1-week) and late (6-week) timepoints. At 1-week post-surgery time point, where we found broad type 2 immune stimulation, there was a significant increase in expression of genes associated with muscle satellite cell activation (*Pax7, Myod1, Myf5, Myog, Mymk*, and *Areg*) and myofiber fusion and development (*Myh3, Myh8*, and *Myl2*) with rSEA treatment compared to saline treated controls (**Fig. 2a, fig. S12e**). The increased expression of muscle development genes with rSEA at early time points correlated with increased muscle tissue. At 6-weeks post-rSEA treatment, immunofluorescent staining of dystrophin, a marker of mature muscle tissue, increased with rSEA treatment compared to saline controls (**Fig. 2b**). Moreover, the rSEA treated muscle had centrally located nuclei, a characteristic of regenerating muscle tissue, compared to peripheral nuclei in the saline treated controls. In addition to increased muscle tissue, rSEA treatment reduced fibrosis. Masson’s trichrome staining of the muscle injury showed reduced collagen deposition without granuloma formation with rSEA treatment (**Fig. 2b**). Gene expression of the muscle tissue at 6-weeks post-injury, supported the morphological findings with decreased expression of fibrosis-associated genes (*Col1a1, Col5a1, Col6a5*), decreased ratio of *Col1a1* to *Col3a1*, and decreased *Tgfb2* with rSEA treatment relative to saline controls (**Fig. 2c**).

**Fig. 2.**
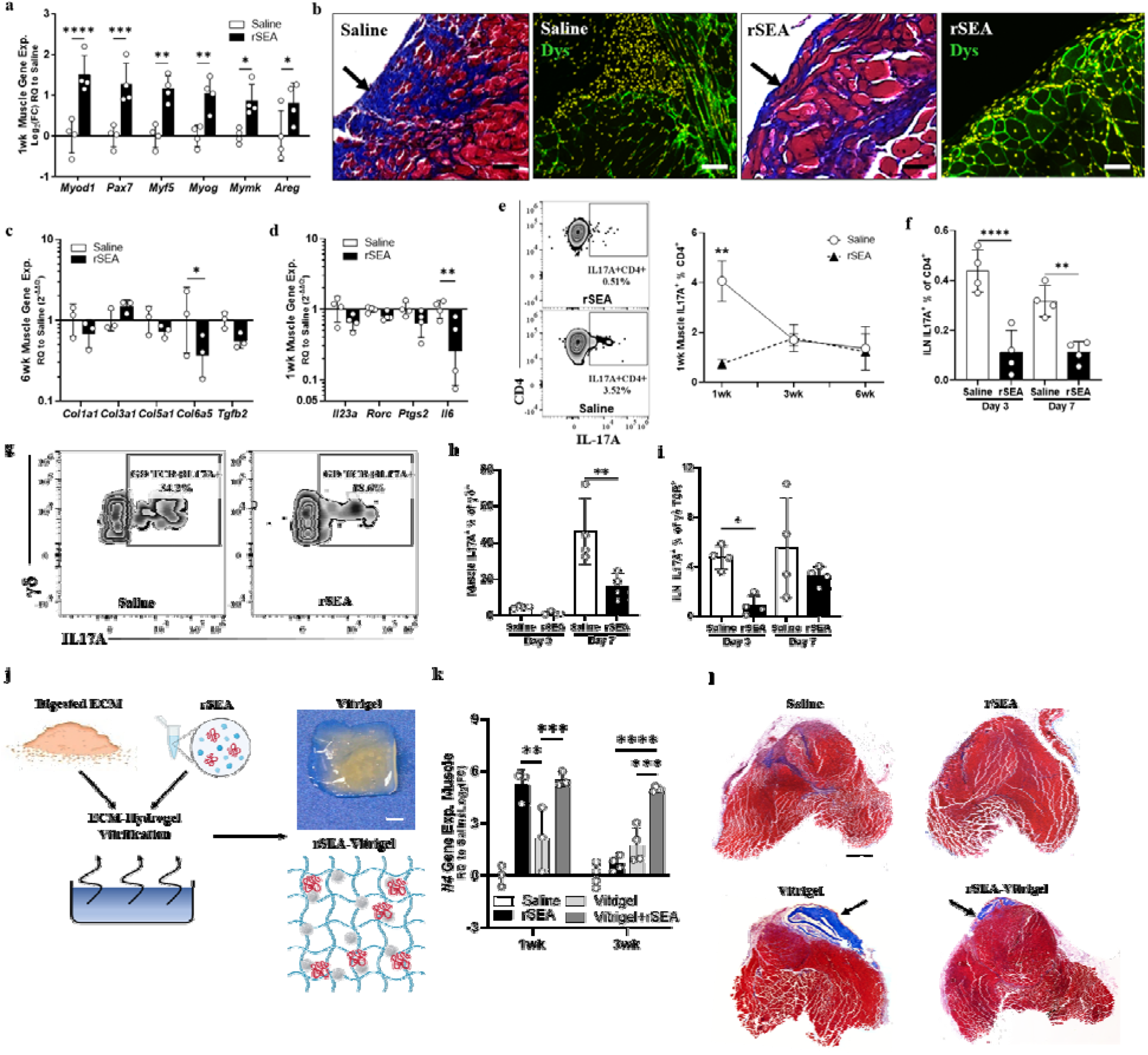
rSEA promotes skeletal muscle repair and decreases fibrosis and type 3 immune responses. **a**. 1-week post-injury expression of genes involved in muscle regeneration in mice treated with either saline or rSEA. **b**. 6-week post-injury muscle histology treated with either saline or rSEA, stained with Masson’s trichrome and immunofluorescent staining (nuclei = DAPI (yellow), mature muscle = dystrophin (green). Arrow indicates site of original muscle resection. **c**. Gene expression of various fibrosis-associated collagens and *Tgfb2* at 6-weeks post-injury and treatment with saline or rSEA. **d**. Gene expression of T_H_17 associated genes at 1-week post-injury and treatment with saline or rSEA. **e**. Representative plot of ICS for IL-17A among muscle CD4^+^ T cells at 1-week (left) and % of CD4 cells expressing IL-17A at 1-, 3- and 6-week (right) after injury and treatment with saline or rSEA as determined by ICS. **f**. % of CD4 cells expressing IL-17A by ICS in iLN 1-week post-muscle-injury and treatment with saline or rSEA. **g**. Representative plot of ICS for IL-17A among muscle γδ T cells at 1-week post-injury. **h**. % of γδ T cells expressing IL-17A in muscle at 1-week post-injury and treatment. **i**. % of γδ T cell expressing IL-17A in iLN at 1-week after injury and treatment with saline or rSEA. **j**. Production summary and gross image of vitrified SIS-ECM combined with rSEA. (Continued on next page.) (Continued from previous page.) **k**. Gene expression of *Il4* in muscle for 1-, 3-, and 6-weeks post-injury and treatment with either saline, rSEA, pure vitrigel or vitrigel formulated with rSEA. **l**. Transverse histological sections of muscles stained with Masson’s trichrome 6-weeks post-injury and treatment with either saline, pure vitrigel and vitrigel formulated with rSEA. Statistical tests represent all biological replicates, and all experiments were replicated at least twice. Graphs show mean ± s.d. (a, c-f, h, i, k), n = 3-4. *\*P* < 0.05, *\*\*P* < 0.01, *\*\*\*P* < 0.001, *\*\*\*\*P* < 0.0001 by two-way ANOVA with Sidak’s multiple comparisons. Scale bars: 100 μm (b), 1 cm (j), and 1 mm (l).

### rSEA treatment decreases IL-17A producing CD4^+^ and γδ^+^ T cells

Schistosome eggs deposited in tissues can induce fibrosis over time and induce granuloma formation similar to the foreign body response. Furthermore, SEA has been used as a model for fibrosis when coated on glass, -sepharose, and -polystyrene beads in multiple tissues including liver and lung (*28, 42*). As fibrosis and granuloma formation are not desirable outcomes in tissue repair, we therefore sought to further evaluate immunological features associated with fibrosis, specifically type 3 (17) immune cells (*43, 44*). Expression of type 3 immune-associated genes including *Il23a, Rorc, Ptgs2 (COX2*) and *Il6* decreased in the muscle tissue 1-week after injury and rSEA treatment (**Fig. 2d**). IL-17A, central to type 3 immune responses, is implicated in fibrosis in multiple tissues including lung, liver, skeletal muscle, and in fibrosis associated with the foreign body response (*43-46*). In parallel with the reduced fibrosis we observed histologically, rSEA treatment significantly decreased IL-17A producing T_H_17 cell number and percentages in the muscle at day 7 post-injury compared to saline returning to similar levels at 3 and 6 weeks (**Fig. 2e**). Further exploring the earlier time points, rSEA induced the most significant decrease in T_H_17 cells at 3 days with a significant decrease still present at 7 days post treatment compared to saline (**Fig. 2f**). rSEA treatment also impacted IL-17A expression by γδ T cells (**Fig. 2g-i)**. While γδ T cell numbers significantly increased after rSEA treatment at 1-week (**fig. S13**), expression of IL-17A^+^γδ^+^ T cells was minimal at 3 days in both groups and significantly decreased with rSEA treatment at 7 days compared to saline treated controls (**Fig. 2g, h**). In the draining inguinal lymph node, IL-17A^+^γδ^+^ T cell percentage significantly decreased at 3 days with rSEA treatment and moderately decreased at 7 days compared to saline control (**Fig. 2i, fig. S13**). We also found a significant reduction in the number of IL17A^+^CD4^+^ cells (and percentage of CD4 and CD45) of rSEA-treated mice at 1-week post-injury, suggesting that decreased inflammatory signatures are occurring more broadly in regionally and systemically in secondary lymphoid tissues (**fig. S14**). Further analysis of fibrosis-related gene expression in the muscle found significant decreases in *Il17a* (signal below threshold in rSEA-treated), as well as genes encoding the canonical T_H_17 transcription factor, *Rorc*, and the primary T_H_17-promoting cytokine, *Il23a* in rSEA treated samples compared to saline treated controls (**Fig. 2d**).

### Release of rSEA from a vitrified ECM hydrogel enhances the type 2 regenerative response

Larger tissue defects may require a scaffold to enable cell migration and the repair of larger tissue volumes. Biological scaffolds derived from tissue ECM, are used clinically for wound healing and reconstruction applications. Preclinical and clinical studies demonstrate that ECM materials can induce type 2 immune responses and promote tissue repair (*37, 47, 48*). Although we see the effects of a single dose of rSEA for weeks, the ability to incorporate rSEA into a biomaterial would provide a scaffold for larger tissue defects and deliver the immunotherapy over an extended period of time.

Biological scaffolds are available in the form of powders and sheets but ECM hydrogels that can encapsulate drugs are generally weak and quickly dissolve (*38*). To create more robust ECM hydrogels that can release proteins and lipids in a controlled manner, we applied a vitrification process that evaporates water in a controlled humidity and temperature so that the macromolecular assembly can occur while a drug or biologic is encapsulated (**Fig. 2j, fig. S15a**). Vitrification of ECM gels derived from urinary bladder ECM increased matrix assembly and fibril formation, similar to previous studies with collagen gels (*49*) as visualized by gross histology and transmission electron microscopy (TEM) (**fig. S15b**). The ECM hydrogel water content decreased after vitrification in parallel with an increase in mechanical properties (G’ modulus) further confirming ECM assembly and formation of a stronger material (**fig. S15c**). Release profile of a model small molecule drug from the vitrified ECM confirmed controlled release over 7 days (**fig. S15d**).

To evaluate the biological activity of the controlled release formulation, we applied particles of vitrified ECM, with or without encapsulated rSEA, to the murine VML. A single dose of rSEA and the vitrigel-encapsulated rSEA stimulated similar levels *Il4* in the muscle tissue at one week, however only the Vitrigel+rSEA extended the increased *Il4* expression to the 3-week time point compared to a single dose of rSEA with a 30-fold increase in *Il4* expression (**Fig. 2k**). The vitrified ECM hydrogel alone stimulated *Il4* gene expression to a small degree. Histological analysis further supported enhanced muscle healing with Vitrigel+rSEA treatment without fibrosis (**Fig. 2l**). The Vitrigel ECM provided a scaffold for tissue growth in addition to controlled release of the rSEA that resulted in a larger volume of new muscle tissue. Vitrigels without rSEA are still visible in the muscle after 6-weeks compared to vitrigels with rSEA that were largely degraded, likely due to increased repair. Moreover, the overall volume of new muscle tissue was smaller in the vitrigel alone group.

### rSEA promotes healing in articular cartilage and cornea tissue injury models

A type 2 immune response and IL-4 expression is associated with repair in multiple tissues beyond muscle including liver, articular cartilage, the central nervous system, and skin (*21, 26, 41, 50-52*). To determine if rSEA could be broadly applied to promote regeneration in tissues beyond muscle, we evaluated the therapeutic potential of rSEA in cartilage and cornea injury models. For cartilage repair, we used the anterior cruciate ligament transection (ACLT) model that induces articular damage, loss of cartilage and development of osteoarthritis (*53*). We injected rSEA intra-articular (IA) two and three weeks after the ACLT injury (**Fig. 3a**) and evaluated the joints 4 weeks after injury compared to vehicle injections.

**Fig. 3.**
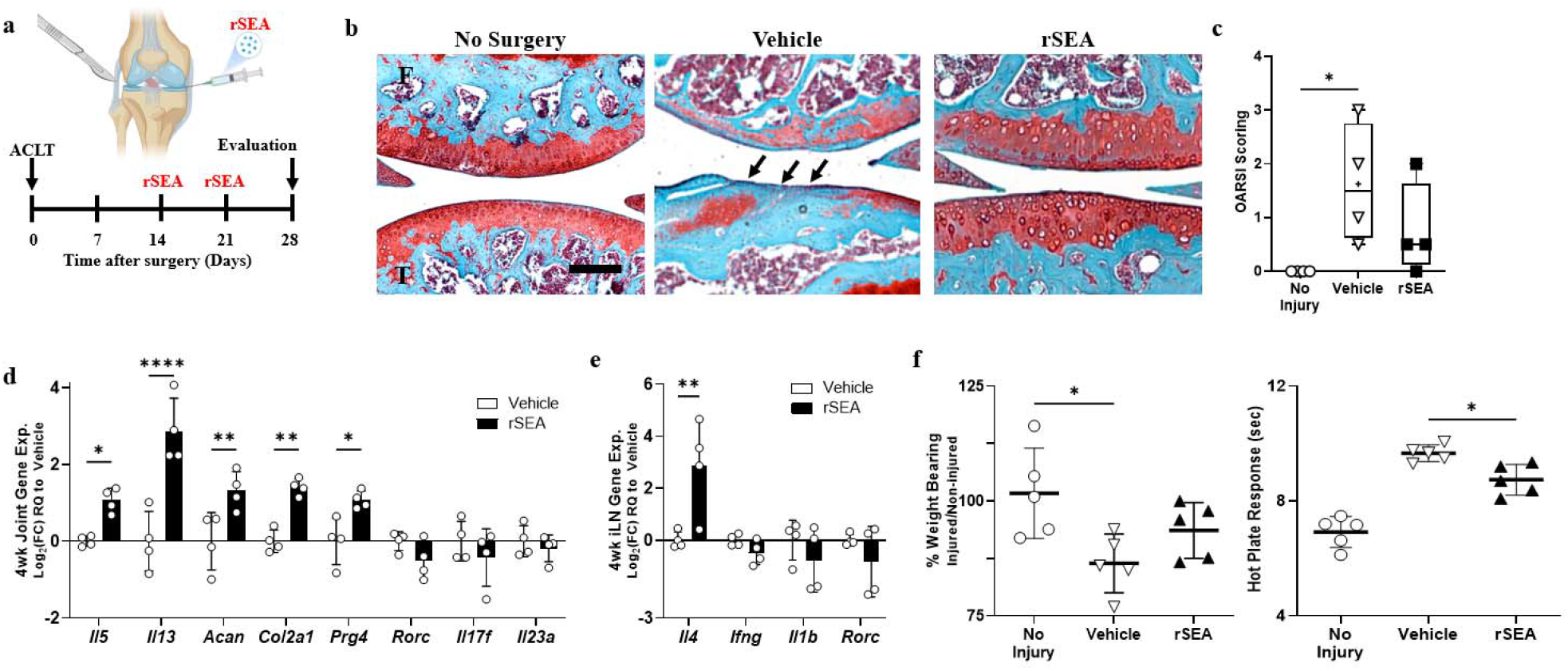
rSEA immunotherapy promotes repair in articular joint injuries. **a**, ACLT injury model timeline. **b**, Representative images from Safranin-O stains of uninjured joints (no surgery) or injured joints treated with vehicle or rSEA 4 wks post-ACLT. Arrows indicate where cartilage damage has occurred. **c**, OARSI scoring of uninjured joints or injured joints treated with vehicle or rSEA 4 wks post-ACLT. **d**, Articular joint gene expression of type 2, type 3 immune genes and cartilage extracellular matrix genes Aggrecan (*Acan*), Type 2 collagen (*Col2a1*), and Lubricin (*Prg4*) 4-weeks post-ACLT, treated with vehicle or rSEA. **e**, iLN gene expression of type 2 and type 3 immune genes 4-weeks post-ACLT. **f**, Hot plate reaction times and weight bearing assessments in mice without injury or 4-weeks post-ACLT treated with vehicle (saline) or rSEA. Statistical tests represent all biological replicates, and all experiments were replicated at least twice. Graphs show mean ± s.d. (c-e), n = 4-5, box and whisker plot with median as central line, and ‘+’ designates mean (c). *\*P* < 0.05, *\*\*P* < 0.01, *\*\*\*P* < 0.001, *\*\*\*\*P* < 0.0001 by one-way ANOVA with Tukey’s multiple comparisons (c, f), two-way ANOVA with Sidak’s multiple comparisons (d, e). Scale bars: 100 μm (b).

Histological assessment of the articular joint structure and cartilage using Safranin-O staining for proteoglycans found that rSEA resulted in higher proteoglycan staining in the cartilage layer, improved tissue structure, and trended in higher levels of repair quality as measured by the semi-quantitative OARSI scoring system compared to vehicle controls (**Fig. 3b, c**). Gene expression analysis of the whole joint tissue at the 4-week time point supported a type 2 immune skewing. Expression of type 2 genes (*Il5* and *Il13*) and cartilage repair and extracellular matrix markers (*Prg4, Acan1*, and *Col2a1*) increased compared to vehicle controls (**Fig. 3c**). Similar to the muscle injury after rSEA treatment, we found that immune type 1 and type 17 gene expression signatures (*Rorc, Il17f*, and *Il23a*) trended downward in comparison to vehicle controls (**Fig. 3c**). To evaluate functional repair, we tested nociception and weight bearing of the injured limb. Hotplate nociception measures the latency period of hindlimb response to heat-induced pain, wherein shorter time responses indicate lower inflammation and more repair (*53*). We found that rSEA treatment significantly decreased the nociceptive response time of the mice in comparison to vehicle treatment suggesting reduced pain (**Fig. 3d**). Functional weight bearing analysis, a measure of the percentage of weight placed upon the injured limb relative to the uninjured limb at improved with rSEA treatment (Vehicle: 86.44 %, rSEA: 93.61 %) (**Fig. 3d**).

We further evaluated therapeutic potential of rSEA in a cornea wound that is similarly characterized by poor healing capacity and scar formation when damaged. In the corneal debridement injury model, resident stromal cells are activated leading to fibrosis and scarring which results in limited vision (**Fig. 4a**) (*54, 55*). We injected rSEA in the subconjunctival space immediately after injury and compared the tissue response to control saline injections. Gross imaging of the corneal surface and tissue clarity 2-weeks after injury showed a significant increase in cornea repair with rSEA treatment compared to controls as measured by blinded quantitative analysis of the scar area (**Fig. 4b**). Immunofluorescence staining of alpha smooth muscle actin (αSMA), a marker that correlates with fibrosis-associated corneal vascularization, also decreased with rSEA treatment compared to control wounds further suggesting improved healing (**Fig. 4c**). We assessed gene expression of the injured and rSEA-treated corneas and found a significant decrease in inflammatory and angiogenic associated genes 1-week post-treatment including *Lyve1, Cd31, Vegfc, Cd36*, and *Acta2* (**Fig. 4d**).

**Fig 4.**
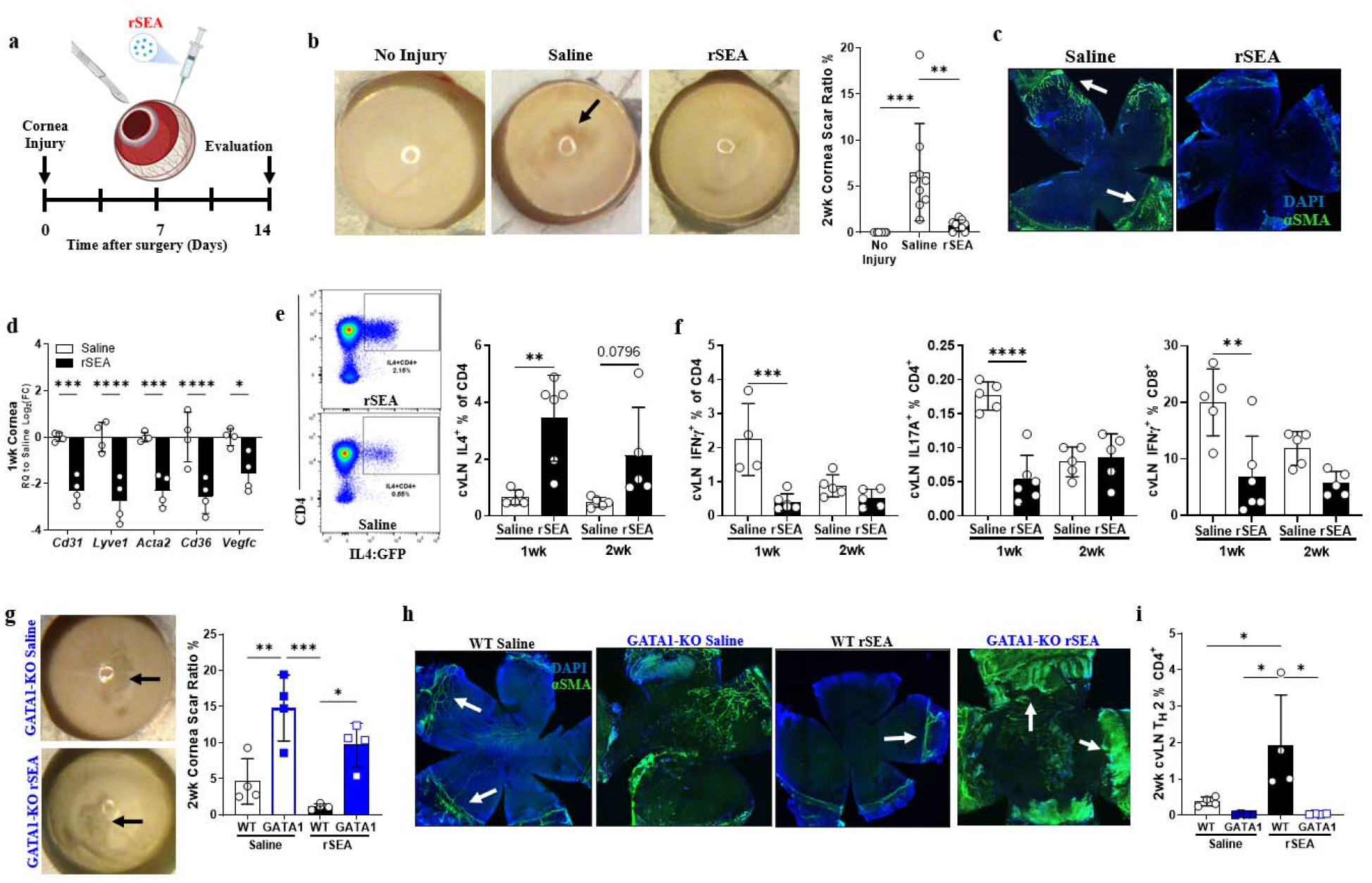
rSEA immunotherapy promotes repair in corneal injury. **a**, Cornea debridement injury model timeline. **b**, Corneal gross images and scar ratio assessment at 2-weeks post-injury treated with saline vehicle or rSEA, two study replications combined. **c**, Immunofluorescent images of nuclei (DAPI-blue) and α-SMA (green) staining on wounded corneas, 2-weeks post-injury and treated with saline or rSEA. **d**, Corneal expression of genes associated with scar vascularization at 1-week post-injury and treatment with saline or rSEA. **e**, Representative flow plots of 2-week post-injury 4get cervical LNs (cvLN) and % T_H_2 populations at 1-week vs. 2-weeks post-injury and treatment with saline or rSEA. **f**, ICS from cvLN of T_H_1, T_H_17, and IFNg^+^CD8^+^ T cells (%) at 1-week & 2-weeks post injury and treatment with saline or rSEA. **g**, Representative gross images of wounded corneas from GATA1 KO mice treated 2-weeks with saline or rSEA and scar area ratio assessment. **h**, Immunofluorescent staining of nuclei (DAPI, blue) and -SMA (green) on corneas 2-weeks post-injury and treatment with saline or rSEA in WT vs. GATA1 KO mice. **i**, cvLN ICS for IL-4 in WT vs. GATA1 KO mice 2-weeks post-injury and treatment with saline or rSEA. Statistical tests represent all biological replicates, and all experiments were replicated at least twice. Graphs show mean ± s.d. (b, d-f, g, i), n = 4-6. *\*P* < 0.05, *\*\*P* < 0.01, *\*\*\*P* < 0.001, *\*\*\*\*P* < 0.0001 by One-way ANOVA with Tukey’s multiple comparisons (b), and two-way ANOVA with Sidak’s multiple comparisons (d-f, g, i).

We then examined the immune response in the cornea after wounding and rSEA treatment to see if an increased type 2 profile correlated with increased tissue repair similar to the muscle and cartilage. We observed few changes in the immune cell populations in the cornea tissue as measured by flow cytometry which may be due to the small cell numbers even when multiple corneas are combined (**fig. S16**). In the draining cervical lymph nodes however, there was a significant increase in IL-4 -expressing CD4^+^ T cells one week after rSEA treatment and moderate increase at 2 weeks (**Fig. 4e**). In parallel, there was a significant decrease in IL-17A^+^CD4^+^, IFN-γ^+^CD4^+^, IFN-γ^+^CD8^+^ T cells percentage with rSEA treatment 1-week post-treatment with minimal changes at 2 weeks (**Fig. 4f**).

Since rSEA treatment increased eosinophil migration in other tissues we hypothesized that eosinophils may be contributing to rSEA-mediated cornea repair. In the ΔdblGATA model that does not have eosinophils, the scar area significantly increased in size with or without rSEA treatment (**Fig. 4g**). Furthermore, the αSMA immunofluorescence staining increased with or without treatment suggesting impaired wound healing and increased fibrosis (**Fig. 4h**). Immunological analysis of the draining cervical lymph node revealed that the significant increase in T_H_2 cells induced by rSEA in WT animals is completely ablated in the ΔdblGATA (**Fig. 4i**). This finding contrasts with the VML model (**Fig. 1e**) and thus suggests that eosinophils are important effectors in healing of the cornea wound by rSEA-induced T_H_2 responses. Thus, rSEA treatment acts on multiple cell types, whose importance in the regenerative process depends on the tissue type and wound.

## Discussion

In this work we designed a pro-regenerative immunotherapy derived from fractionated helminth parasite egg antigens and demonstrate their ability to enhance wound healing and deter fibrosis post-traumatic injury across three injury models. Taking the eggs from *S. mansoni* helminths, we derived an alternative formulation from the soluble egg antigen, rSEA. We showed that the rSEA stimulated a type 2 immune signature in lymphoid cells and myeloid cells, further decreasing pro-inflammatory immune polarization, and later timepoints revealed decreased levels of fibrosis associated with inhibition of T_H_17 and γδ^+^IL-17A^+^ cells. Application of rSEA to muscle, cornea, and articular joint injuries generally improved tissue healing assessed by gene expression signatures, cell populations, and/or histological assessment. Controlled release of rSEA from a natural sourced and decellularized biomaterial hydrogel further promoted healing and regeneration of larger tissue volumes. Our rSEA formulation, particularly in the form of an ECM hydrogel, is therefore a regenerative immunotherapy with potentially broad application to tissue repair and homeostasis, though several vital questions remain for exploration in future work such as the optimal formulations and scaling of SEA to benefit healing, deleterious off-target effects, and immunotherapy-induced susceptibility to other pathogens during treatment.

A type 2 immune response is central to how the immune system responds to helminth infection. While the type 2 response has long been considered an anti-helminth response, it is also now hypothesized that helminths may induce this anti-inflammatory immune signature to repair the damage caused to the host, thereby enhancing mutual survival. Recent studies highlight the importance of the context of expression of type 2 associated molecules that are important in dictating outcomes, such as Amphiregulin, IL-13 and IL-33 (*20*). IL-4 and type 2 immunity is associated with tissue repair and healing in multiple tissue types including liver (*50*), bone (*51*), cartilage (*48*), muscle (*4, 5*), corneal (*55*), and nervous tissues (*26, 52*), suggesting that the rSEA may be broadly applicable for tissue repair (*18, 20, 21*). Treatment with rSEA induced an immune profile that included eosinophils and T_H_2 cells producing significantly higher levels of IL-4, IL-5, and IL-13 protein or gene expression compared to injured tissue without treatment. The type of tissue where injury and treatment may impact which cells are responding to rSEA and promoting tissue repair. In our studies, application of rSEA to a cornea wound in the ΔdblGATA murine cornea injury model completely abolished repair. However, in skeletal muscle injuries the absence of eosinophils in ΔdblGATA mice did not completely ablate pro-healing gene expression signatures induced by rSEA despite their well-recognized role in muscle tissue repair (*5*). This suggests that rSEA may activate multiple immune cell populations to promote repair that differ according to tissue type. It is also likely that rSEA influences stromal, stem, or progenitor cell populations in addition to immune cells as Helminth infections were shown to stimulate stem cells in the intestinal niche (*19*).

As type 2 immune responses have also been implicated in long-term fibrotic and allergic responses (*21*), further studies may elucidate how these immune responses differ from the regenerative response. Chronic helminth exposure results in fibrosis and is postulated to be from an overzealous wound-healing response (*18, 21*). The mechanical properties of worm casings and the secretions from the eggs together may also contribute to helminth-related fibrosis as we observed with biomaterial hydrogels with increasing stiffness that produced more fibrosis in part due to IL-17 signaling (*42*). Recently, it was found that *S. mansoni* eggshells alone can temporarily deter the foreign body response and that the antigens secreted through the eggshell induce granulomas to control the timing of egg extrusion into the environment to continue the parasite’s lifecycle (*42*). In our regenerative studies with the rSEA therapy, we did not observe fibrosis and instead found a reduction in IL-17 and fibrosis markers.

While we purified SEA to make the regenerative formulation, it is still a complex mixture of proteins, proteoglycans and lipids. Multiple components of SEA have been identified and the known functions of these individual components support their potential in promoting tissue repair. It is likely that the different components of rSEA target different and complementary cellular functions in the immune and stromal compartments of a wound and tissue to stimulate type 2 immune responses and tissue repair. Further isolation of SEA components may thus reduce therapeutic response. ECM-derived biomaterials are an example of a complex mixture where a mixture of components contribute to the therapeutic efficacy. Multiple ECM-derived materials are FDA-approved as medical devices and biologics depending on the source and clinical application. Manufacturing protocols and biological release assays enable consistent products that have a strong track record of safety and efficacy. As *in vitro* culture of *Schistosomiasis* has already been demonstrated (*56*), reproducible culture and scale up may be possible. Taken together, the development of rSEA and the tissue repair findings describe a regenerative medicine approach that targets the immune system and presents a new class of immunotherapies with potential broad application across multiple tissue and organ systems.

## Acknowledgments

We thank the Biomedical Research Institute (BRI) of Rockville, Maryland, for providing helminth eggs and SEA associated resources, contract: HHSN272201700014I, and Christopher A. Moad for technical assistance. Thank you to Edward J. Pearce for his critical review of the manuscript.

## Funding

National Institutes of Health Pioneer Award DP1AR076959 (JHE)

Bloomberg∼Kimmel Institute (JHE, DMP)

Morton Goldberg Professorship (JHE)

The Department of Defense, Award Number W81XWH-19-1-0576 (JHE)

The National Eye Institute grant, R01EY029055 (JHE)

The National Science Foundation Graduate Research Fellowship Program #DGE-1746891 (DRM, ANP)

The Armed Forces Institute for Regenerative Medicine II award number WFUHS 441081 CF-11 (AEA)

The NIH T32 Training Grant 1T32AG058527-01 and (JIA)

The NIH T32 Training Grant 5T32CA153952-08 (JIA)

The NIH K-Award 1KL2TROO1429 (EM)

The Rhines Rising Star Larry Hench Professorship (EM)

## Author contributions

Conceptualization: DRM, JHE, and DMP

Data Curation: KK, DRM

Investigation: DRM, LC, JIA, JSTH, ANP, EM, JCM, XW, JH, HZ, AFC, HHN, YC

Formal Analysis: DRM, JH, JSTH, XW, LC

Funding acquisition: DRM, JHE, DMP

Project administration: JHE, DMP, DRM

Supervision: JHE, DMP

Validation: DRM, LC

Visualization: DRM, HHN

Writing – original draft: DRM, JHE, DMP

Writing – review & editing: All authors

## Data and materials availability

All relevant analyzed data present in this manuscript, Figures or the associated Supplemental Figures, Supplementary Materials will be made available to peer reviewers upon request. Raw and post-processed data sets for flow cytometry and qRT-PCR will be annotated and made publicly available in accordance with MIFlowCyt and MIQE recommended guidelines upon formal publication of this study. A dedicated effort was made to adhere to conventional standards for all studies herein to ensure reproducibility. All *S. mansoni* eggs, livers, and reagents utilized in this work were obtained courtesy of the Biomedical Research Institute (BRI) (Rockville, Maryland, USA), a 501 (c)(3) nonprofit research company supported by the National Institute of Allergy and Infectious Diseases (NIAID) of the National Institutes of Health (NIH) through the NIH-NIAID, Contract: HHSN272201700014I.

